# Mayaro virus replication restriction and induction of muscular inflammation in mice are dependent on age and type-I interferon response

**DOI:** 10.1101/602920

**Authors:** CM Figueiredo, RL Neris, DG Leopoldino, JS Almeida, JS dos-Santos, CP Figueiredo, M Bellio, MT Bozza, I Assunção-Miranda

## Abstract

Mayaro virus (MAYV) is an emergent Arbovirus belonging to the Alphavirus genus from the *Togaviridae* family which has been circulated in forest regions of American continent through small outbreaks. Recent studies warned for the risk of MAYV dispersion to new areas and for the potential establishment of an urban epidemic cycle. Similar to Chikungunya and other arthritogenic Alphavirus, MAYV-induced disease shows a high prevalence of arthralgia and myalgia that can persist for months. Despite this, knowledge regarding pathogenesis, characteristics of host immune response, and resolution of MAYV infections are still limited. Here we investigated the dependence of age, innate and adaptive immunity for the control of MAYV replication and induction of inflammation in mice. We observed that age and type I interferon response are related to restriction of MAYV infection and tissue inflammation in mice. Moreover, we showed that MAYV continues to replicate persistently in adult recombination activation gene-1 efficient mice (RAG1^−/−^), indicating that adaptive immunity is essential to MAYV clearance. Despite chronic replication, infected adult RAG1^−/−^ mice did not develop an apparent signal of muscle damage at late infection. On the other hand, MAYV infection induces muscular and paw inflammation in young WT and adult Type I Interferon receptor deficient mice (IFNAR^−/−^). In addition, MAYV infection triggers an increase in the expression of pro-inflammatory mediators, such as TNF, IL-6, KC, IL-1β, MCP-1, and RANTES, in muscle tissue, and decreases TGF-β expression. Taken together, our study contributes to the comprehension of MAYV pathogenesis, and describes a translational mouse model for further studies of MAYV infection, as well for testing vaccine and therapeutic strategies against this virus.

**Author Summary:** MAYV-induced disease presents a high prevalence of arthralgia and myalgia that potentially persist for months, which is characteristic of the arthritogenic Alphavirus group. However, information regarding MAYV infection and the molecular mechanism of pathogenesis is still scarce. Here we investigated the dependence of age, innate and adaptive immunity for the control of MAYV replication and induction of inflammation in mice. We observed that tissue inflammation and the restriction of MAYV replication in mice are affected by aging and type I interferon response. Besides, we also showed that adaptive immunity was important for MAYV clearance in adult mice. Histological analyses demonstrated that MAYV replication triggered muscular and paw inflammation in young WT and adult type-I interferon receptor deficient mice. In addition, the level of expression of several pro-inflammatory cytokines was increased in the muscle MAYV-infected mice. Our data provide an advance for understanding the molecular mechanism involved in MAYV pathogenesis, as well as describes an *in vivo* model for further investigations on MAYV infection and for antiviral compounds and vaccine testing.

## Introduction

Mayaro virus (MAYV) is an *Alphavirus* from the Togaviridae family, transmitted to humans mainly by the bites of *Haemagogus* mosquitoes (1). MAYV was first isolated in 1954 from a febrile case in Trinidad and Tobago and maintained until the present day on restricted circulation in Central and South American forest regions on sporadic outbreaks (2–4). However, recent studies indicate that the number of reported MAYV cases could be underestimated, warning for the risk of emergence, dispersion to new areas, and for the potential establishment of an urban epidemic cycle (1, 5–9). Even in face of such risks, information regarding MAYV infection and mainly the molecular mechanism of pathogenesis is still very limited.

Due to the profile of clinical manifestations, MAYV is grouped with the arthritogenic Alphavirus such as Chikungunya (CHIKV) and Ross River (RRV). MAYV infection promotes a febrile condition that presents a set of unspecific signs and symptoms, such as rash, headache, and ocular pain, which facilitates its misdiagnosis as other arboviroses such as dengue fever (2, 10–12). Moreover, MAYV infected patients present a high incidence of articular and muscular pain (2, 7), reaching about 50% and 77% of patients in some outbreaks, respectively (12). In addition, it has also been reported that myalgia and articular symptoms of MAYV infections could persist for months, revealing a common feature to arthritogenic alphavirus-induced disease (13–16)].

High activation of immune response has been described in CHIKV and RRV-infected patients presenting acute and persistent symptoms (17, 18). Analyses of muscle biopsies of CHIKV-infected patients with severe polyarthralgia and myalgia showed that symptoms persistence was associated with long-term cellular infiltrate at articular and muscle tissue (19). However, the characteristics of the immune response induced by MAYV, the mechanisms of resolution of the infection or symptoms persistence are largely unknown. The one-year longitudinal study of Santiago et al. 2015 demonstrated that MAYV-infected patients also present prolonged immune response, with high concentrations of pro-inflammatory mediators in their serum (20). They found lower amounts of GM-CSF, IL-5, and IL-10 in MYV-infected patients when compared to CHIKV patients, which indicates differences in the profile of the induced immune response. Consistent with this, a difference in cytokine expression between MAYV and CHIKV infection in human U937 cell lineage (21) was observed. However, contrastingly from what was observed in patients, MAYV infected U937 cells display a more anti-inflammatory profile of immune activation. Despite the divergence, this data reinforces the necessity of further studies that evaluate cellular and molecular aspects of MAYV infection.

The muscle and joint inflammation during CHIKV and RRV have been evaluated in immunocompetent and immunodeficient mice, as well as in non-human primates [(22–24). It was demonstrated that inflammatory monocyte infiltrates trigger tissue damage, contributing to the severity of the disease (25). However, currently, there is no systematic study evaluating replication, tissues damage, and immune activation in MAYV-infected mice. Here we investigated age, innate and adaptive immunity dependence for MAYV replication, and induction of inflammation. We observed that MAYV replication and dissemination in wide-type (WT) SV129 mice is determined by aging and is controlled by innate immunity. We also demonstrated that adaptive immunity is essential to MAYV clearance. Finally, MAYV infection promotes an inflammatory process in the muscle of young WT mice, with high amounts of pro-inflammatory mediators. Taken together, our data contribute to the comprehension of MAYV pathogenesis and offer a mouse model for further studies of MAYV infection and for testing therapeutic strategies against this arbovirosis.

## Methods

### Virus

Mayaro virus (ATCC VR 66, strain TR 4675) was propagated in BHK-21 cells cultured in α-MEM (alfa-Minimum Essential Medium - Invitrogen) supplemented with 10% fetal bovine serum (FBS). BHK-21 cells were infected in a multiplicity of infection (MOI) of 0.1 and 30 hours post infection (hpi) culture medium was collected and centrifuged at 2,000 x g for 10 min to remove cell debris. The clarified medium was aliquoted and stored at −80 °C. Viral titer of the stock was determined by plaque assay.

### Animals and infection

The experiments were performed using young WT SV129 mice (6, 11, 21 days after birth); as well as adult (8 weeks old) WT SV129, type I interferon receptor deficient mice (IFNAR^−/−^) and WT C57BL/6, recombination activation gene RAG-1 deficient mice (RAG1^−/−^) in the C57BL/6 background. Mice of the desired age were subcutaneously inoculated in the left footpad with 10^6^ pfu of MAYV, using a final volume of 20 μL. Only for the infection of IFNAR^−/−^ mice the inoculation was that of 10^5^ pfu of MAYV. The same volume of uninfected BHK-21 medium (Mock) was used as the control. Each experimental group was housed individually in polypropylene cages with free access to chow and water. Young mice were housed with the uninfected mother during all the experiment.

Mice were weighed daily and clinical signals were scored. The area of hind limb foot edema in IFNAR^−/−^ animals was determined from the width-height measurements of the metatarsal region using a digital caliper. Tissue samples were collected at 2 and 4 days post infection and stored at −80 °C until processed or fixed in 4% formaldehyde. All experimental procedures performed were approved by the Institutional Animal Care and Use Committee of the Federal University of Rio de Janeiro (protocol no. 014/16).

### Virus quantification

MAYV titer and viral load in tissue samples were determined by plaque assay in BHK-21. Tissue samples were homogenized in α-MEM using a fixed relation of mass/volume and a serial dilution was prepared in α-MEM (ten-fold). Then, each dilution was used to infect confluent BHK-21 cells seeded in 24-well plates. After 1 hour of adsorption, the medium was removed and 2 mL of 1% carboxymethylcellulose (w/v) (Sigma-Aldrich) in α-MEM supplemented with 2% FBS were added onto the infected monolayer, and then cells were incubated at 37 °C. After 48 h, cells were fixed using 4% of formaldehyde and plaques were then visualized by staining the fixed monolayer with 1% crystal violet in 20% ethanol. A title was calculated as plaque forming units per mL (pfu/mL) and in tissue samples converted to pfu/mg.

### Histology

The muscle and footpad were collected at defined days post infection and fixed with 4% of formaldehyde for 24 hours. The footpad was decalcified using EDTA solution (125g/L, pH 7.0) and then fixed. Tissues were embedded in paraffin after dehydration. Paraffin-embedded tissue sections of 5 μm were prepared and stained with hematoxylin and eosin (H&E). Images were obtained using optical microscopy with a magnification of 10 X (Olympus BX40), and images were acquired using software Leica Application Suite 3.8 (Leica).

### Cytokine quantification by qPCR

Hind limb muscles were homogenized in DMEM using a fixed relation of 0.2 mg of tissue/μL, and 200 μL of the homogenate was used for RNA extraction with Trizol (Invitrogen) according to the manufacturer’s instructions. Purity and integrity of RNA were determined by the 260/280 and 260/230 nm absorbance ratios. One μg of isolated RNA was submitted to DNAse treatment (Ambion, Thermo Fisher Scientific Inc.) and then reverse-transcribed using the High-Capacity cDNA Reverse Transcription Kit (Thermo Fisher Scientific Inc). Quantification of cytokines expression was performed using the Power SYBR kit (Applied Biosystems; Foster City, CA). Actin was used as an endogenous control. Primer sequences were the following: IL-6 (FW-5′-TCA TAT CTT CAA CCA AGA GGTA-3′; REV, 5′-CAG TGA GGA ATG TCC ACA AAC TG-3′), IL-10 (FW, 5′-TAA GGG TTA CTT GGG TTG CCA AG-3′; REV, 5′-CAA ATG CTC CTT GAT TTC TGG GC-3), TGF-β (FW, 5′-GAC CGC AAC AAC GCC ATC TA-3′; REV, 5′-AGC CCT GTA TTC CGT CTC CTT-3′), MCP-1 (FW, 5′-GTC CCC AGC TCA AGG AGT AT-3′; REV, 5′-CCT ACT TCT TCT CTG GGT TG-3′), KC (FW, 5′-CAC CTC AAG AAC ATC CAG AGC-3′; REV, 5′-AGG TGC CAT GAG AGC AGT CT-3′), TNF (FW, 5′-CCT CAC ACT CAG ATC ATC TTC TCA-3′’; REV, 5′-TGG TTG TCT TTG AGA TCC ATG C-3′), IL-1β (FW, 5′-GTA ATG AAA GAC GGC ACA CC-3′; REV, 5′-ATT AGA AAC AGT CCA GCC CA-3′); Actin (Forward: 5’-TGT GAC GTT GAC ATC CGT AAA-3’ and Reverse: 5’-GTA CTT GCG CTC AGG AGG AG-3’).

### Statistical analyses

The statistical analyses were performed comparing means by two-tailed T-Student’s tests using Graph Pad Prism version 7.00 for Windows, Graph Pad Software, La Jolla California USA, www.graphpad.com.

## Results

### MAYV inoculation induces clinical signs of infection in young and type-I interferon receptor deficient mice

Clinical findings suggested that MAYV induces a robust inflammatory response in patients, similar to other arthritogenic alphavirus (20). However, MAYV tissue tropism and damage induced by infections has not been characterized yet. Hence, we first characterized whether MAYV infects or impacts animals in an age- and immunological status-dependent manner. For this, MAYV was inoculated in left hind limb footpad of young (6, 11, 21 day-old) and adult WT SV129 mice. The infection resulted in high lethality and severe weight loss for young animals (≤11-days-old) (**Fig 1A and 1B**). Interestingly, 21-day-old infected-mice had no change in body weight when compared with Mock-infected mice and were completely resistant to MAYV-induced lethality, indicating their ability to control MAYV replication.

**Fig 1.**
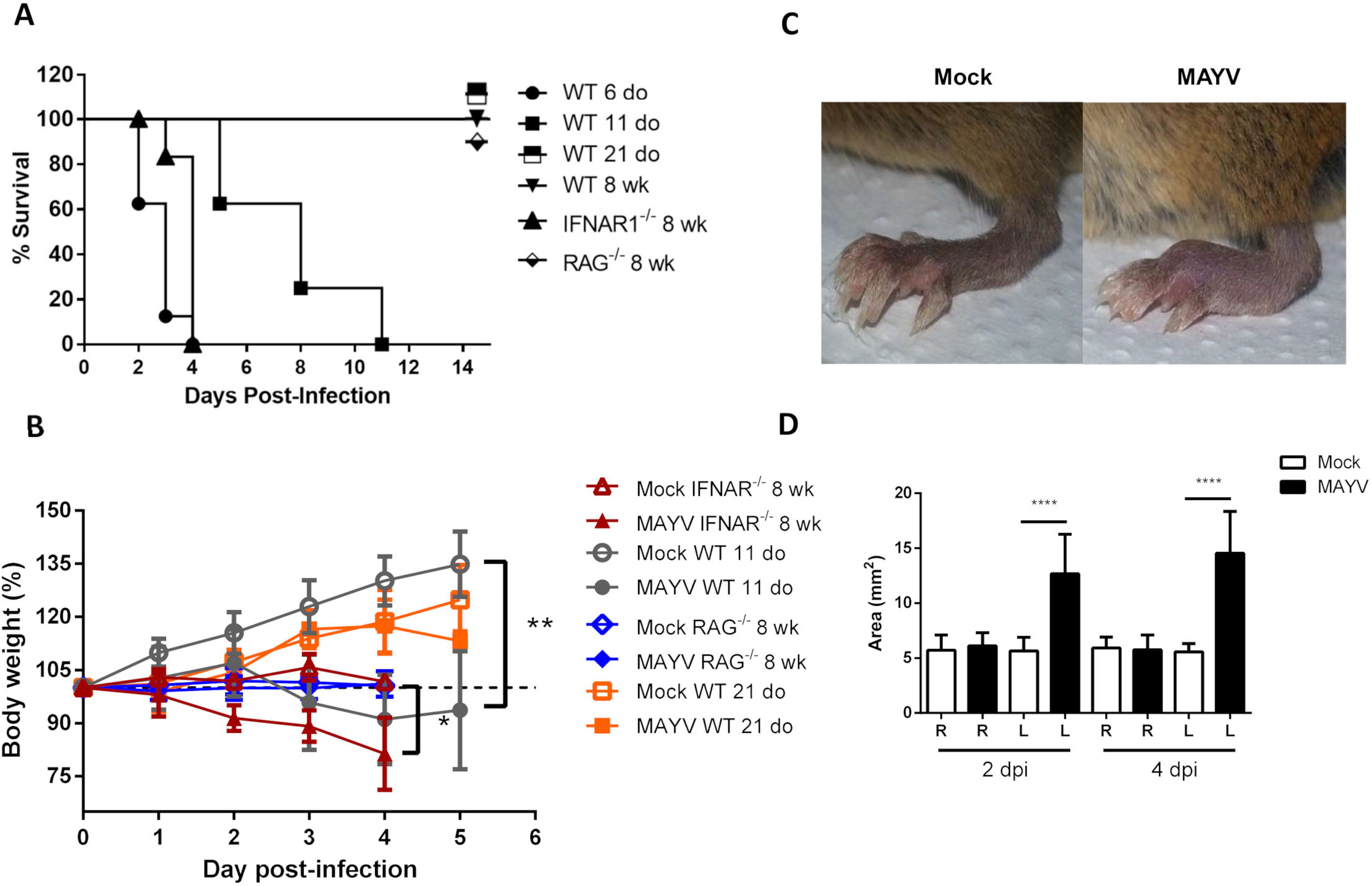
MAYV infection induces clinical signs in young WT and type I interferon receptor deficient mice. WT SV129 mice of different ages (do, days old), 8 week-old (wk) SVA129 (IFNAR^−/−^) and C57BL/6 RAG1^−/−^ mice were subcutaneously infected with MAYV in the left footpad. **(A)** Survival was monitored up to 14 days post infection for mice, and **(B)** body weight up to 5 days post infection. **(C)** Representative image of MAYV or Mock infected left footpad in IFNAR^−/−^ mice. **(D)** The swelling area of left (L) and Right (R) paws was obtained using the measurement of paw height and width obtained by a digital caliper, at 2and 4 days post infection. Values were plotted as mean ± Standard Deviation (SD). Statistical analyses were performed comparing means by two-tailed t-student. *□0.05; **□0.01;**** □0.0001.

Inoculation of MAYV on adult WT SV129 and C57BL/6 mice did not cause any clinical signal of infection when compared to mock infected animals (**Table 1 and S1A fig**). However, inoculation of MAYV in type-I interferon receptor deficient (IFNAR^−/−^) adult mice results in early lethality and severe weight loss, similarly to infection of 6-day-old WT mice (**Fig 1A and 1B**). In addition, we observed intense paw edema in MAYV-injected footpad, when compared with mock (**Fig 1C**), or UV-inactivated MAYV infected footpad (data not shown). Measurement of edema area showed that the swelling in IFNAR^−/−^ was present at day 2 post infection (dpi) and maintained until 4 dpi (**Fig 1D**). No alteration was observed in the counter lateral footpad. In addition to the swelling, IFNAR^−/−^ mice presented lethargy, locomotion dysfunction, and posterior weakness. Similar diversity and intensity of clinic signals were observed in WT 11-day-old mice (**Table 1**).

**Table 1.** Clinical signs in MAYV infected mice.

|  | Mice |  |  |  |  |  |  |
| --- | --- | --- | --- | --- | --- | --- | --- |
| Clinical Signs | IFNAR<sup>-/-</sup><br>(SVA129)<br>8 wk | WT<br>(SV129)<br>6 do | WT<br>(SV129)<br>11 do | WT<br>(SV129)<br>21 do | WT<br>(SV129)<br>8 wk | WT<br>(C56BL/6)<br>8 wk | RAG1<sup>-/-</sup><br>(C57BL/6)<br>8 wk |
| Lethargy | +++ | +++ | +++ | + | N | N | N |
| Weight loss | ++ | - | ++++ | + | N | N | N |
| Locomotion dysfunction | + | - | ++ | N | N | N | N |
| Posterior weakness | ++ | ++ | ++ | N | N | N | N |
| Posterior paralysis | N | + | N | N | N | N | N |

Clinical signs were observed in different day-old (do) or week-old (wk) MAYV-infected mice. (+) Indicates presence and the amount of + the intensity of the signal. (-) Indicates that it was not possible to evaluate and (N) Absence of the signal.

We also investigated MAYV infection in adult recombination activation gene-1 deficient mice (RAG1^−/−^), which do not maturate B and T lymphocytes (26), used as a model to investigate the involvement of adaptive immunity in diseases (25). MAYV infection of adult RAG1^−/−^ mice did not cause lethality, weight loss, and even other clinical signals observed in young WT and adult IFNAR^−/−^ mice (**Table 1**). Together, these results indicate that young mice and IFNAR^−/−^ are highly susceptible to MAYV infection, while adult mice are resistant even in the absence of lymphocytes.

### MAYV replication restriction is dependent on innate and adaptive immunity

Subsequently, we performed a temporal analysis of MAYV viremia to correlate clinical signals with the ability to control viral burden. We observed an increase in the amount of MAYV infectious particles in the blood of mice of all ages on the first day of infection (**Fig 2A**). However, blood viral load was sustained at about 10^6^ pfu until 4 dpi in 11-day-old WT mice, while continuously decreased in 21 day-old WT mice until undetectable levels at 5 dpi, which reinforces the age-dependence for the control of MAYV infection. In agreement with this, the infection of WT SV129 and C57BL/6 adult mice results in a brief viremia. This ability to control viral replication was completely lost in the absence of type I interferon response, as observed by the continuous increase of viremia until the lethality of IFNAR^−/−^ mice (**Fig 2A**). Despite the absence of clinical signs, MAYV infectious particle titers were sustained for a long period and maintained at about 10^4^-10^5^ pfu/ml until 7 dpi in the blood of RAG1^−/−^ mice. Conversely, in this same period, MAYV was no longer detected in the blood of adult WT C57BL/6.

**Fig 2.**
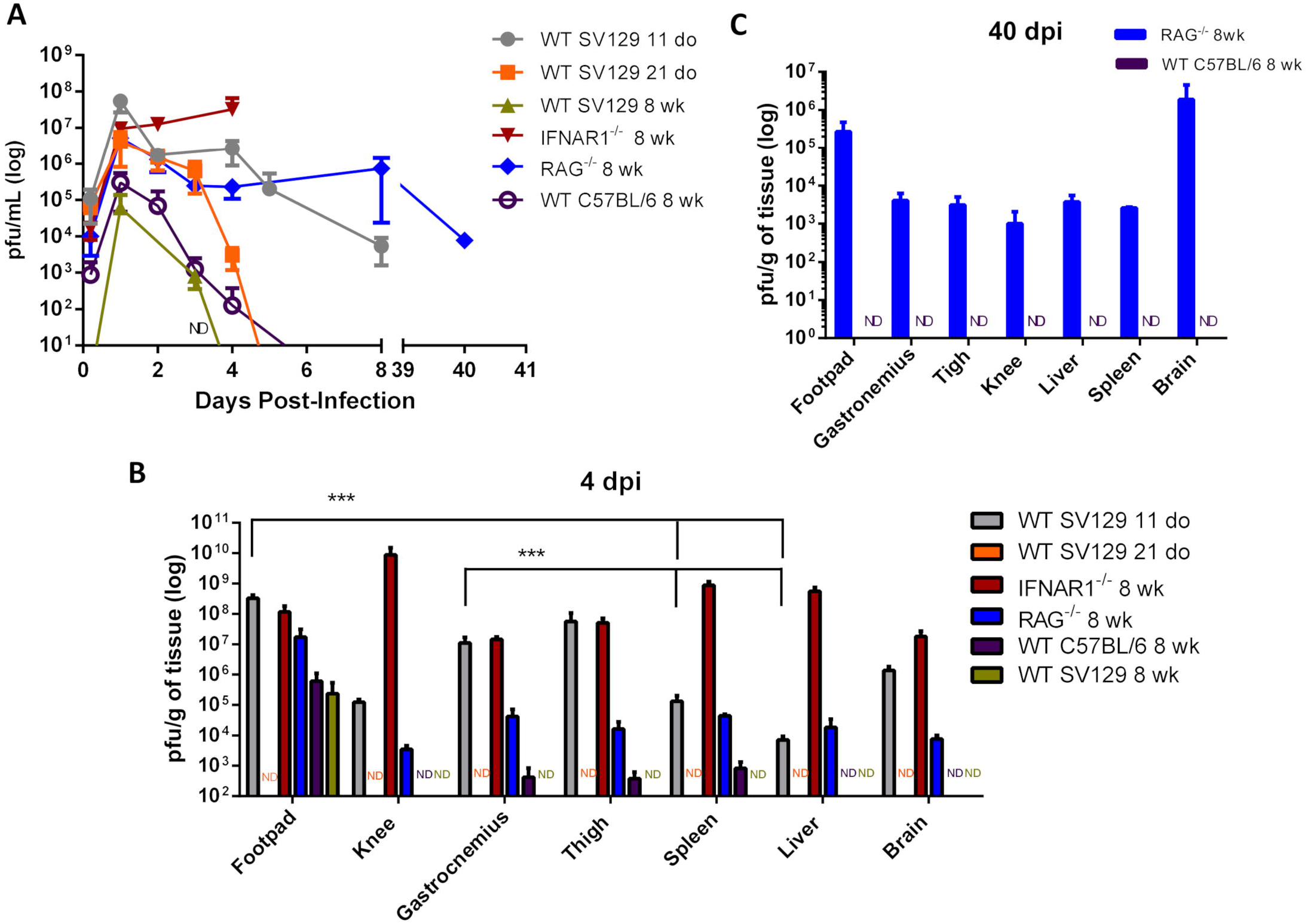
Control of viremia and tissue viral loads is dependent on age, innate and adaptive immunity. Mice were infected with MAYV in the left footpad. **(A)** Temporal analyses of MAYV infectious particles present in mice blood were determined by plaque assay. **(B)** MAYV load at different tissues 4 days post infection; **(C)** 40 days post infection. Tissue samples were homogenized using a fixed relation of weight/volume and titled by plaque assay. Values were plotted as mean ± Standard Deviation (SD).Statistical analysis on tissues within each group was performed by two-tailed t-student. * □0.05 and ** □0.01.

In order to identify preferential areas of replication, we evaluated the tissue distribution of MAYV at 4 dpi. MAYV infectious particles were detected in the articular and muscular tissues, as well as in liver, spleen, and brain of 11-day-old WT and adult IFNAR^−/−^ mice (**Fig 2B**). Although MAYV has a widespread distribution in mice tissues, we could observe a preferential distribution to the paw and the skeletal muscle in 11-day-old WT mice, since they presented a significantly higher viral load when compared to other peripheral tissue, as liver and spleen (**Fig 2B**). A high viral load was also detected in muscular and articular tissues from the opposite side of the injection site (**S1B fig**). These broad distributions were not observed in the infection of the 21-day-old and 8-week-old WT animals.

### MAYV establishes persistent infection in the absence of adaptive immunity

Although adult RAG1^−/−^ mice presented a lower viral load in all tissues when compared to young WT and adult IFNAR^−/−^ mice 4 dpi, MAYV load was higher in RAG1^−/−^ mice than in WT C57BL/6 mice and viral dissemination seems to be more efficient since it was detected in the brain tissue of RAG1^−/−^ mice (**Fig 2B**). In addition, analysis of the blood and tissues of RAG1^−/−^ mice demonstrated that MAYV continues to replicate actively in these animals until 40 days post infection, despite no apparent morbidity (**Fig 2A and 2C**). These results indicate that adaptive immunity is determinant for the elimination of MAYV, thus contributing to avoid chronicle infection. In addition, the high viral load at 40 dpi was found in the left and right foot, and in the mice’s brains (**Fig 2C and S1C Fig**), suggesting the existence of preferential reservoir sites of MAYV for the maintenance of persistent infections.

### MAYV induces inflammation and muscular damage in young WT and adult IFNAR^−/−^ mice

Since joint and muscular tissues were the main sites of MAYV replication in 11-day-old mice, we investigated whether virus replication triggers pathological alterations. Histological analysis of H&E stained skeletal muscle of the hind limb of young WT and IFNAR^−/−^ mice showed sites of injury, with necrosis, edema, and infiltration of inflammatory cells at 4 dpi with MAYV (**Fig 3**). MAYV-induced muscular damage was similar in young WT and adult IFNAR^−/−^ mice. However, the extension of lesions seems to be higher in young WT mice. Despite the inability to clear the viral infection, no histological alteration was found in muscles of RAG1^−/−^ mice even at 40 dpi (**Fig 3**), correlating with the absence of clinical signals in these mice. This evidence indicates that adaptive immunity activation is determinant for viral clearance and could also be an important factor contributing to MAYV-induced inflammation and lesions.

**Fig 3.**
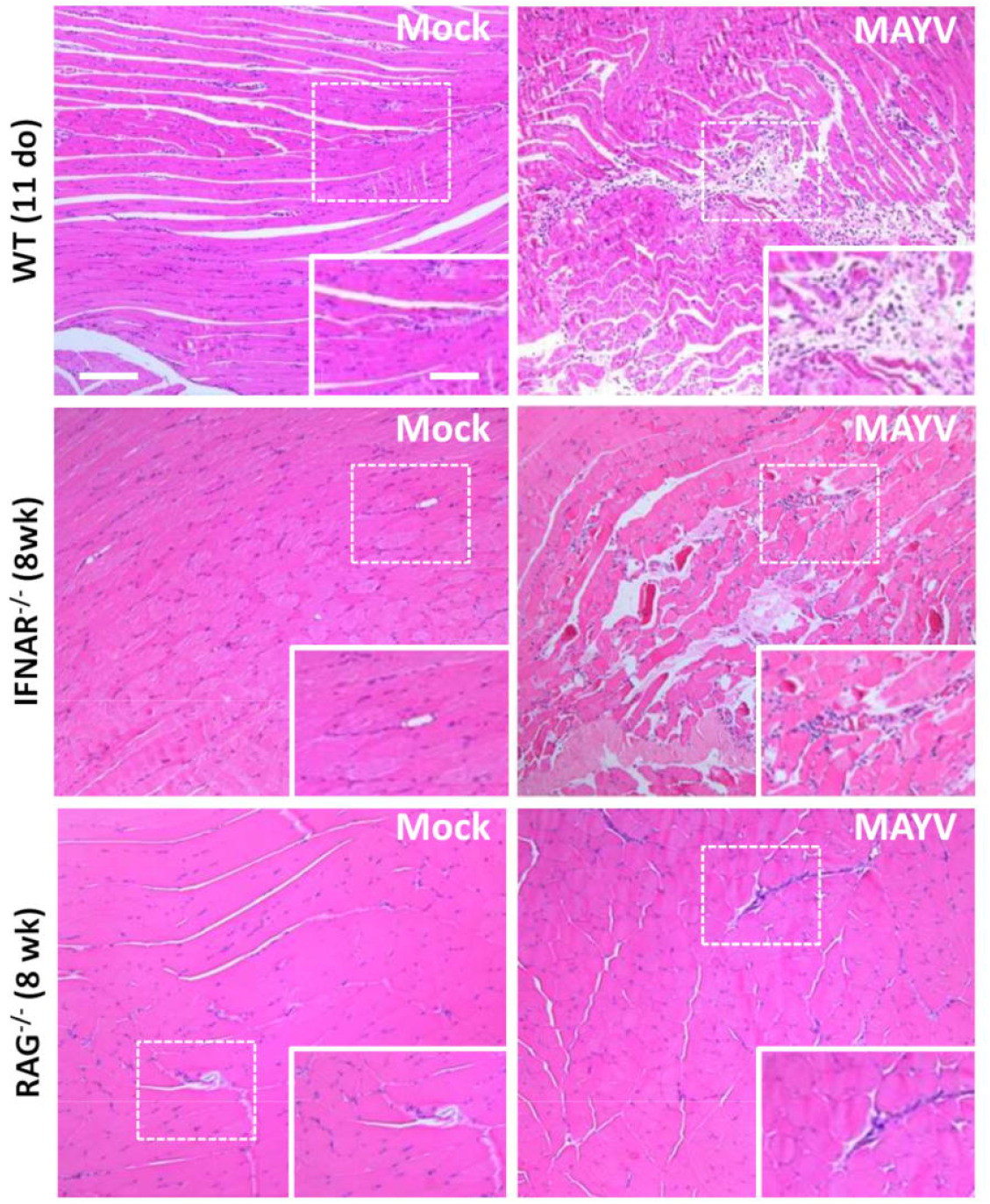
MAYV infection promotes muscle inflammation and necrosis in young WT and IFNAR^−/−^ mice. Eleven day-old (do) WT mice, 8 week-old (wk) IFNAR^−/−^ and RAG1^−/−^ mice were subcutaneously infected with MAYV in the left footpad and muscle tissue was collected and fixed at 4 dpi (WT and IFNAR^−/−^) or 40 dpi (RAG1^−/−^) for histological analysis. Tissues were embedded in paraffin after dehydration and tissue sections of 5 μm were prepared and stained with H&E. Scale bar = 100 μm. Higher magnification images of the regions defined by dashed white rectangles (Scale bar = 10 μm)

Analysis of H&E staining of the hind limb footpad at 4 dpi revealed that MAYV infection resulted in paw inflammation, with an edema area which was close to articular-associated skeletal muscle, mainly in IFNAR^−/−^ infected mice (**Fig 4**). In addition, muscle damage was observed in young WT mice but not in IFNAR^−/−^. Taken together, muscular and paw alterations indicate that MAYV replication in young and adult IFNAR^−/−^ mice results in damage and inflammation in target tissues. The damage induced by infection seems to correlate not only with the viral load, since higher loads of MAYV in the muscle and footpad of IFNAR^−/−^ do not results in higher damage, but might be triggered by inflammatory mediators and cellular activation during the course of infection.

**Fig 4.**
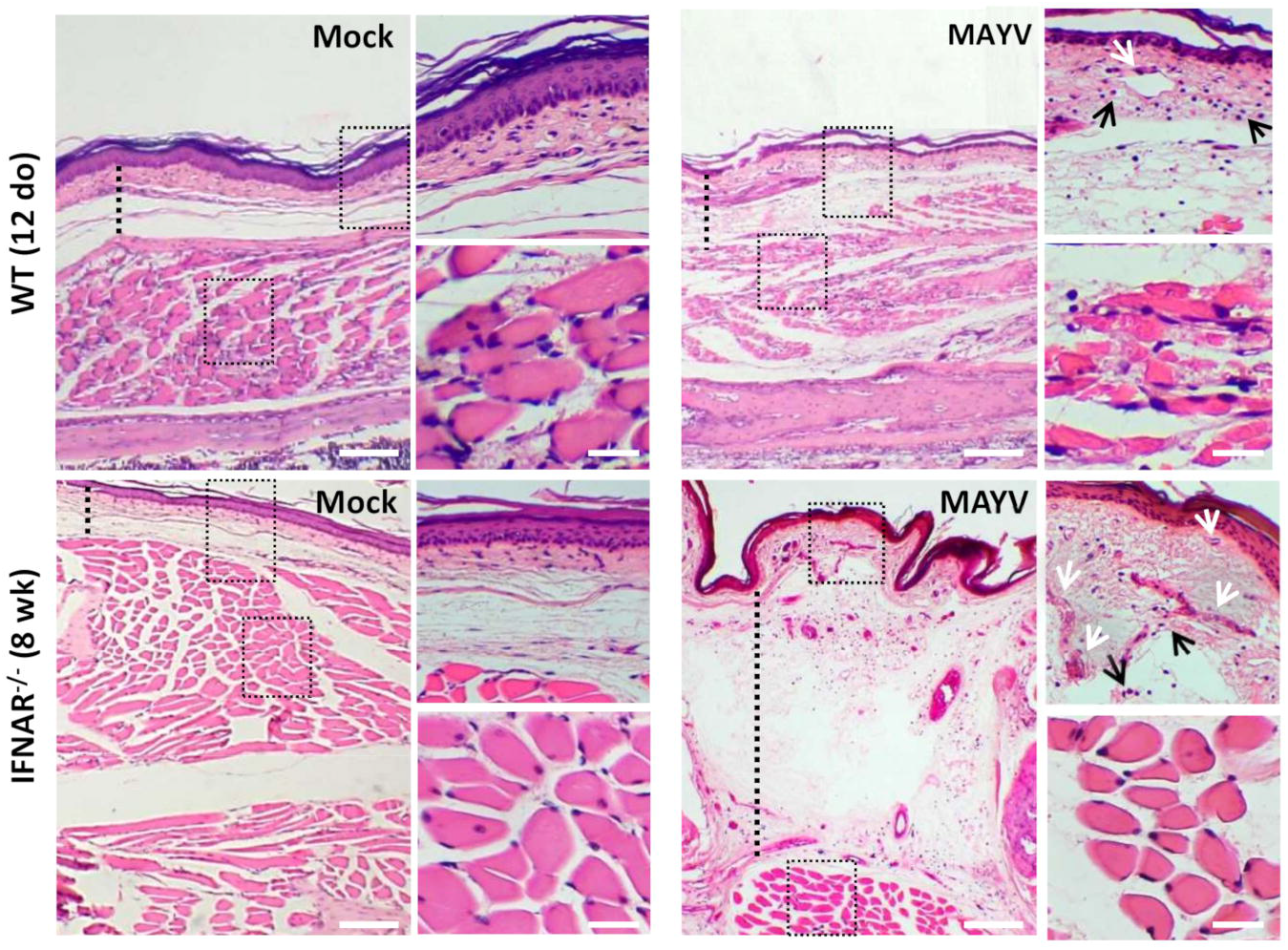
MAYV infection results in inflammation on the paw of young WT and IFNAR^−/−^ mice. 11-day-old (do) WT mice and 8-week-old (wk) IFNAR^−/−^ were subcutaneously infected with MAYV in the left footpad. Footpad longitudinal sections from MAYV- or Mock-infected animals (4 dpi) were stained with hematoxylin and eosin (H&E). Representative images of footpad from MAYV- and Mock-infected animals. Black arrows: inflammatory cell infiltration. White arrows: blood vessels congestion. Scale bar of panoramic images = 500 μm. Higher magnification images of the regions defined by dashed black rectangles (Scale bar = 30 μm).

### MAYV infection triggers a high expression of pro-inflammatory mediator in muscular tissues

Some cytokines have been described as determinant to the progression of CHIKV induced lesions (27). Furthermore, MAYV patients with long-term articular symptoms present high concentrations of pro-inflammatory cytokines in their serum (27). Thus, we investigated if MAYV-induced damage was associated with the induction of a pro-inflammatory response in the muscular tissue of young WT and adult IFNAR^−/−^ mice. The quantification of inflammatory mediator expression by qPCR showed that MAYV replication in muscular tissues triggers high expression of cytokines and chemokines, such as TNF, IL-6, KC, IL-1β, MCP-1, and RANTES (**Fig 5**). The levels of cytokine induction following MAYV infection in IFNAR^−/−^ and young WT were very similar, except for RANTES and KC (**Fig 5**). Since we found that the type-I interferon response was determinant for MAYV infection restriction, we also assessed whether MAYV was able to induce IFN-β expression in young WT mice. We found that MAYV infection promotes an increase of about 15-fold of IFN-β mRNA expression (**Fig 5**). We also evaluated the expression of TGF-β and IL-10, important anti-inflammatory mediators related to a regenerative response. We found that TGF-β was down modulated in young WT mice, while being unaltered in adult IFNAR^−/−^ mice (**Fig 5**). IL-10 expression was not significantly altered in young WT mice and adult IFNAR^−/−^ mice infected with MAYV.

**Fig 5.**
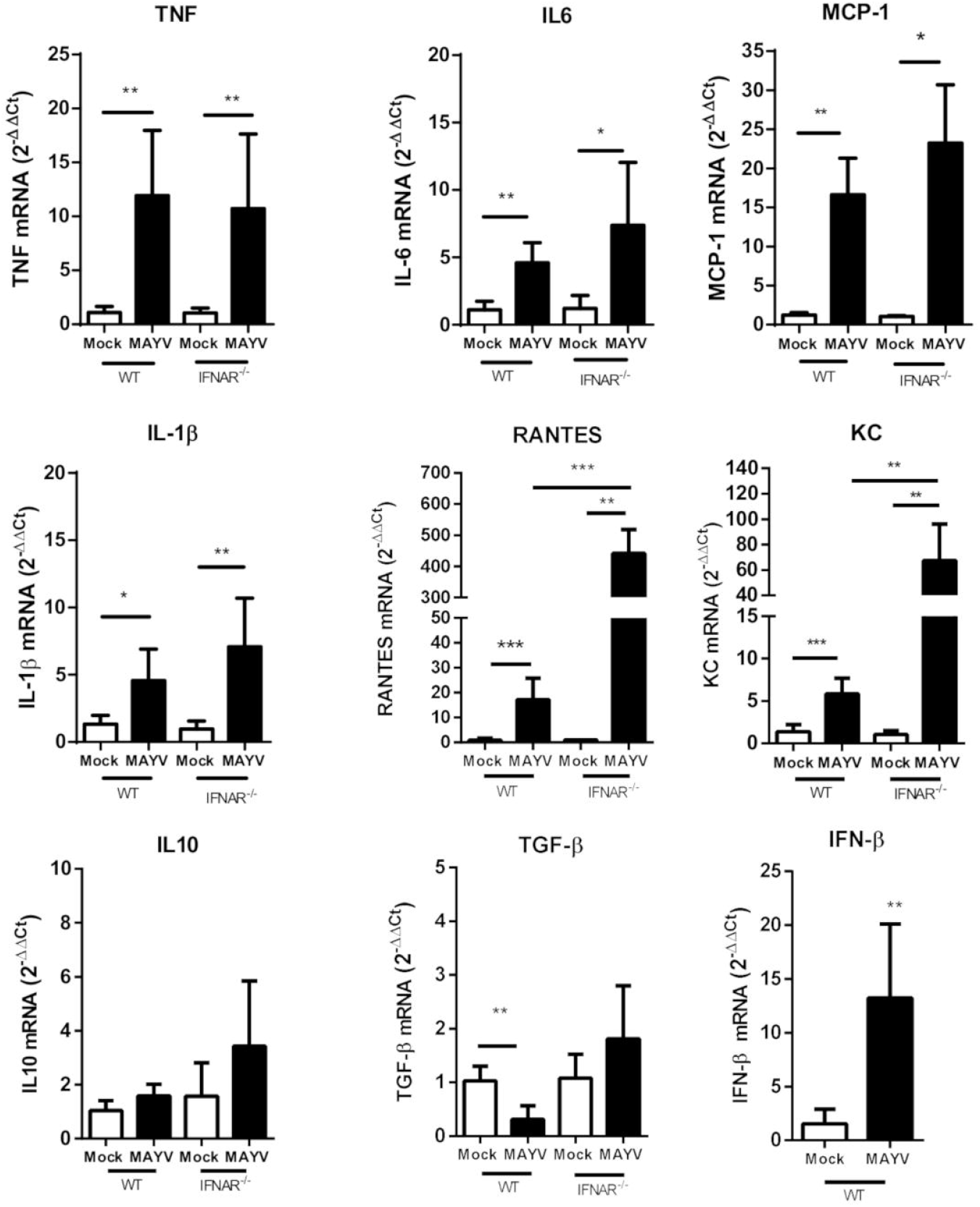
MAYV infection triggers a high expression of pro-inflammatory mediator in muscle tissue. 11-day-old WT and 8-week-old IFNAR^−/−^ mice were subcutaneously infected with MAYV in the left footpad. At four days post-infection, both posterior gastrocnemius muscles were collected and processed to determine the relative expression of pro-inflammatory cytokines. The levels of TNF-α, IL-6, MCP-1, IL-1β, RANTES, KC, IL-10, TGF-β, IFN-β were determined through RT-PCR comparing gene expression with endogenous β-actin expression. Values are showed as mean ± Standard Deviation (SD). Statistical analysis was performed using two-tailed T-student. P *□0.05; **□0.01;*** □0.001.

## Discussion

Host restriction of viral infection could be determined by several aspects of virus-cell interactions, including viral recognition and efficient innate and adaptive immune responses. Here we showed that MAYV replication, induces clinical signals and muscular inflammation in young WT and adult IFNAR^−/−^ mice, which indicates they could be good models to study different aspects of MAYV-induced disease. We found that MAYV presented a very narrow window of time to induce apparent infection in WT mice, with clinical signals been observed in 11-day-old mice but completely unapparent after 21 days of age. The clearance of viremia and absence of MAYV detection at 4 dpi in 21-day-old and adult mice reinforce that mice gain the capability to control virus dissemination with age. Viral replication in young mice could be associated with the inability of the immature immune system of mice to sustain an efficient antiviral response (28, 29). This feature also correlates with clinical evidence of the severity and chronicity prevalence of Alphavirus infection in children and the elderly population (30–32).

The type I interferon (IFN-I) response has been described as essential to controlling RRV, CHIKV, ONNV, and SINV replication (33). The induction of some interferon stimulated genes (ISG) is critical to restrict Alphavirus infection and dissemination (34–36). In agreement with this, adult IFNAR^−/−^ mice infected with MAYV develop a severe and lethal infection, indicating that IFN-I response is also crucial to restrict MAYV infection. However, we observed an increase of IFNβ expression in the muscle of infected young WT mice. It is possible that other factors could be associated with age-dependent severity of the infection or MAYV could inhibit IFNAR signaling in young mice.

Adaptive immunity seems to have a role in the controlling of MAYV infection, mainly in viral clearance. MAYV infection in RAG1^−/−^ mice was completely unapparent and had no histological evidence of muscle damage, even with sustained viremia and significant viral loads in tissues until 40 dpi. MAYV persisted mainly in joint-associated tissue and brain, but infectious particles were detected at lower levels in several peripheral tissues of the mice. The involvement of adaptive immunity in chronicity of CHIKV infection has been demonstrated, but its role in the process of tissue injury could be quite variable in different Alphavirus infection (37, 38). As opposed to MAYV, CHIKV infected RAG1^−/−^ mice presented chronic synovitis and myositis (37, 38). Moreover, RRV induces muscle inflammation and disease in RAG^−/−^ mice during acute infection (39). These data indicate that the role of adaptive immunity in MAYV infection could differ from that of other arthritogenic Alphavirus, an important issue that needs further investigation.

MAYV presented a broad tissue distribution in susceptible mice although in young WT mice the highest load was detected at muscular and articular tissues. In addition, we observed skeletal muscle necrosis and inflammation, mainly in young WT infected mice. The muscular tropism and damage could correlate with the main clinical symptoms observed in MAYV and is characteristic of myositis induced by other arthritogenic Alphavirus (12, 40). Currently, few studies investigated cellular tropism and consequences of MAYV replication (21, 41–43). It was demonstrated that MAYV replicates in RAW macrophages *in vitro*, promoting an increase in the amounts of reactive oxygen species and TNF (43). However, their roles in MAYV pathogenesis are not clear yet. Macrophage activation has a crucial role in articular and muscular inflammation-derived lesions in RRV and CHIKV infection (22, 44). In addition, the production of type I IFN by activated inflammatory monocytes has been suggested as crucial to control acute RRV infection in mice (25). Here, we observed moderate cellular infiltrate on the focus of necrosis and high expression of inflammatory mediators in the muscle of MAYV infected mice, including TNF levels that may contribute to tissue damage. Thus, it is possible that cells from the infiltrate could be responsible for the pro-inflammatory response in the muscle tissue.

Some studies evaluated cytokine profiles correlating with severity and chronicity of CHIKV infection (18, 27, 30, 45). The severity of CHIKV was associated with the levels of IL1-β, IL-6, and RANTES (27), with IL-6 already associated with the persistence of symptoms (18). A similar study in MAYV infected patients was conducted by Santiago et al (2015), showing high levels of TNF, IL-6, IL-8, IL-1Ra, IFN-γ, and others in the serum until 12 months post-acute phase (20). Here we observed that acute MAYV infection promotes elevated expression of inflammatory mediators in the muscle tissue, revealing a translation of our model and MAYV induced disease in patients. Although the cytokine profile was similar to that seen in CHIKV infection, we cannot rule out differences in the amplitude of the inflammatory response. Interestingly, we observed that levels of TGF-β expression were reduced in young WT mice infected with MAYV. High levels of TGF-β were associated with a reduced immune response, persistent viremia, and with joint pathology in CHIKV infection in old mice (46). Therefore, the lower levels of TGF-β in MAYV infection could be important to sustain an immune response that restricts muscle damage. Further studies assessing the role of pro- and anti-inflammatory cytokines in the generation or resolution of muscle lesions are important for exploring the possibility of employing these mediators as new therapeutic targets.

Our study presents the first description of molecular aspects of replication restriction of MAYV, together with demonstration of musculoskeletal inflammation and damage as a consequence of *in vivo* infection. The current knowledge about the mechanism of arthritogenic Alphavirus injury promotion was described using mainly CHIKV and RRV infection in mice models (47, 48). However, the characterization of similarities and specificities of Alphavirus-promoted diseases is determinant for the development of effective vaccines and therapies against this group of viruses. Thus, our study also represents an important contribution as an *in vivo* model for further investigations on MAYV pathogenesis and also to test antiviral compounds and vaccines.

## Acknowledgments

This work was supported by grants from Brazilian funding agencies: Fundação de Amparo à Pesquisa do Estado do Rio de Janeiro (FAPERJ); Conselho Nacional de Desenvolvimento Científico e Tecnológico (CNPq) and Coordenação de Aperfeiçoamento de Pessoal de Nível Superior (CAPES), finance Code 001.

## Supporting information

**Sup 1 fig.**
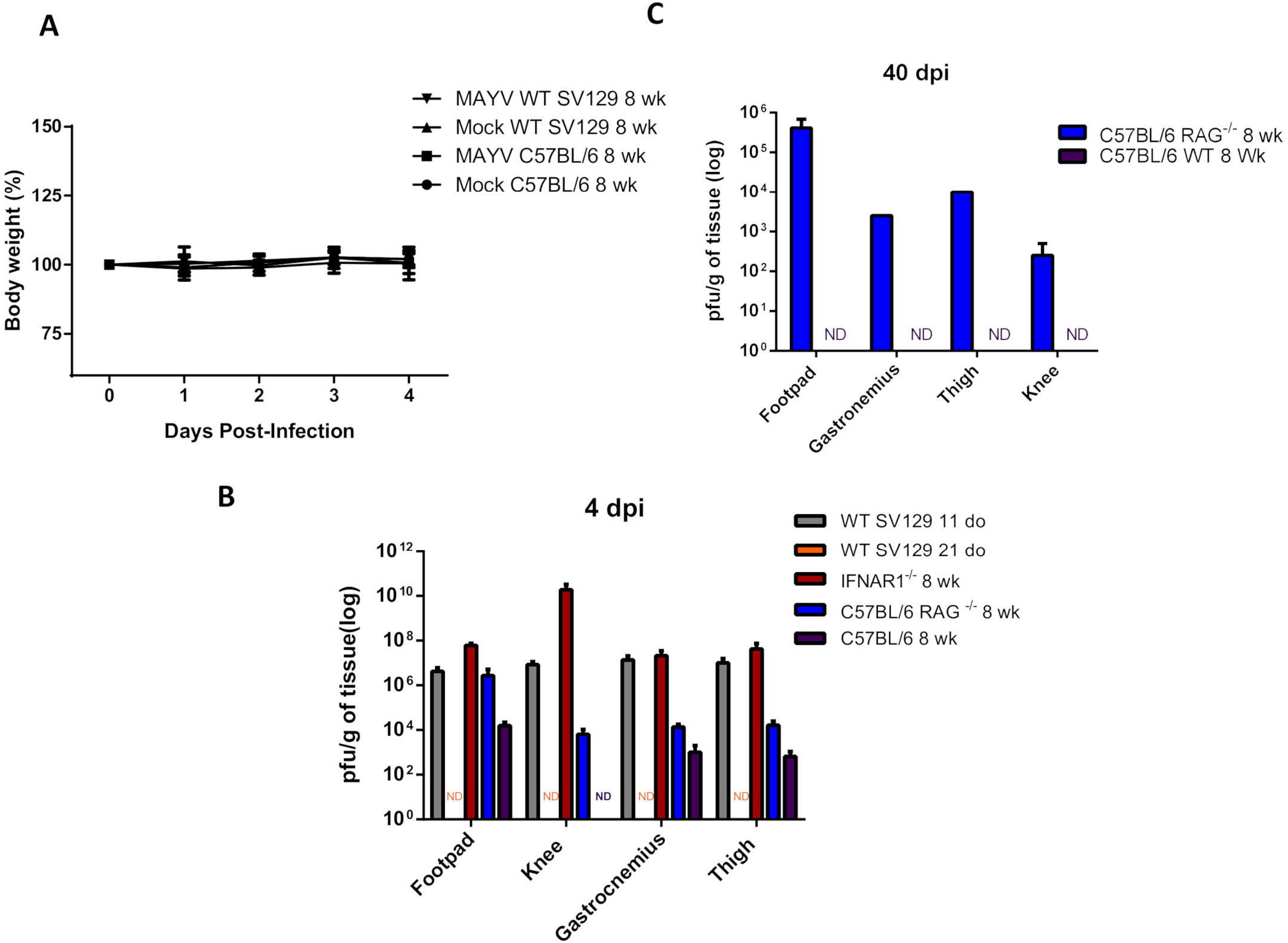
**(A)** Body weight of Adult (8 weeks) Wt SV129 mice and C57BL/6 Wt mice was monitored throughout the days following the infection. Body weight was plotted in % using mass values in the moment of infection as reference. **(B)** MAYV load at right side muscular and articular tissues at 4 dpi and **(C)** 40 dpi in RAG^−/−^ right side tissue. Tissue samples were homogenized using a fixed relation of mass/volume and titled by plaque assay.

